# Molecular architecture of excitatory and inhibitory neurons in the first gustatory relay

**DOI:** 10.64898/2026.07.13.738341

**Authors:** Deepthi Mahishi, Nilay Yapici, Eirene Markenscoff-Papadimitriou

## Abstract

Taste perception is a major determinant of food intake and is dynamically modulated by hunger and satiety signals across species. While the peripheral and cortical representations of taste sensory stimuli are well characterized, far less is known about how metabolic state shapes taste processing in subcortical structures such as the brainstem. The rostral nucleus of the solitary tract (rNTS) is the first central relay for gustatory input, yet how fasting remodels transcriptional programs in rNTS neurons remains poorly defined. To address this gap, we performed cell-type-specific bulk nuclear RNA sequencing of molecularly defined excitatory (*Vglut2^+^*) and inhibitory (*Vgat^+^*) rNTS neurons. We combined Cre-dependent Sun1-sfGFP nuclear tagging with fluorescence-activated nuclear sorting to profile each population in *ad libitum*–fed and 24-hour–fasted mice. The two populations were transcriptionally distinct and differed in their baseline complement of hunger- and metabolic-sensing genes. Fasting elicited a transcriptional response that was largely restricted to excitatory, *Vglut2^+^*, rNTS neurons, while the inhibitory population was minimally affected. In excitatory neurons, fasting reduced the expression of *Gabrd*, the δ subunit of the GABA_A_ receptor that mediates extrasynaptic tonic inhibition, while leaving synaptic GABA_A_ subunits and the glutamatergic or GABAergic neurotransmission machinery unchanged. RNAscope in situ hybridization confirmed *Gabrd* expression in *Vglut2^+^* rNTS neurons and showed that *Gabrd* is also expressed in the non-Vglut2^+^ population; the cell-type specificity of the fasting response therefore reflects differential regulation of a shared gene rather than its restricted expression. Our results identify a cell-type-specific, state-dependent transcriptional signature in which fasting downregulates a mediator of tonic GABAergic inhibition in excitatory rNTS neurons, providing a candidate substrate for the metabolic modulation of taste processing within the brainstem.

**Highlights:**

- Excitatory and inhibitory rNTS neurons are intermingled, not spatially segregated, in rNTS
- Vglut2^+^ and Vgat^+^ rNTS nuclear transcriptomes are distinct at baseline
- Fasting reshapes the Vglut2^+^ transcriptome but leaves Vgat^+^ neurons largely unchanged
- Fasting downregulates *Gabrd*, a mediator of tonic inhibition, in Vglut2^+^ neurons

## Introduction

Taste perception is continuously shaped by animals’ metabolic state: foods that are rejected when sated become acceptable, even rewarding, under conditions of hunger, a phenomenon termed *alliesthesia*^1^. This state-dependent shift in taste perception directly affects food intake, yet the neural mechanisms by which metabolic signals regulate taste processing remain poorly understood. A logical starting point is the first central relay for gustatory information in the brain, the nucleus tractus solitarius (NTS) in the dorsal medulla^2^. NTS is subdivided along its rostro-caudal axis into functionally and anatomically distinct domains that differ in their afferent supply, downstream targets, and behavioral contributions (Figure 1A). The rostral NTS (rNTS) constitutes the first central relay for gustatory and oral somatosensory information, receiving primary afferent input from the chorda tympani and greater superficial petrosal branches of the facial nerve (VII), the lingual-tonsillar branch of the glossopharyngeal nerve (IX), and the superior laryngeal branch of the vagus (X)^3^. The caudal NTS (cNTS), by contrast, is the principal target of vagal viscerosensory afferents conveying mechanical and nutrient-related signals from the gastrointestinal tract. In the mouse, this segregation is evident in coronal section: the rNTS is best sampled at approximately Bregma − 6.72 mm, where the solitary tract lies dorsolateral to a dense field of small-diameter neurons (Figure 1B), whereas the cNTS emerges more caudally at approximately Bregma −7.56 mm, adjacent to the area postrema and dorsal motor nucleus of the vagus (Figure 1C). Taste and oral mechanosensory signals from the anterior tongue and palate are carried by pseudounipolar neurons of the geniculate ganglion, and from the posterior tongue by the petrosal ganglion, with both converging on the rNTS; gut-derived interoceptive signals are carried by nodose ganglion neurons that terminate in the cNTS (Figure 1D). This topographic separation of oral and visceral afferent streams provides the anatomical substrate for the hypothesis that taste and post-ingestive information are differentially gated by internal state at the earliest stage of central processing. Despite the anatomical significance of taste-responsive neurons in the rNTS, most work has focused on thalamocortical processing of taste information in the context of learning and ingestive behaviors^4–7^. Yet the brainstem alone may be sufficient for state-dependent feeding: decerebrate rats, which are devoid of the forebrain but with intact brainstem structures, show state-dependent modulation of feeding behaviors^8^. This underscores the need to investigate the molecular and physiological role of rNTS and brainstem neurons in taste processing and the integrative control of feeding.

**Figure 1.**
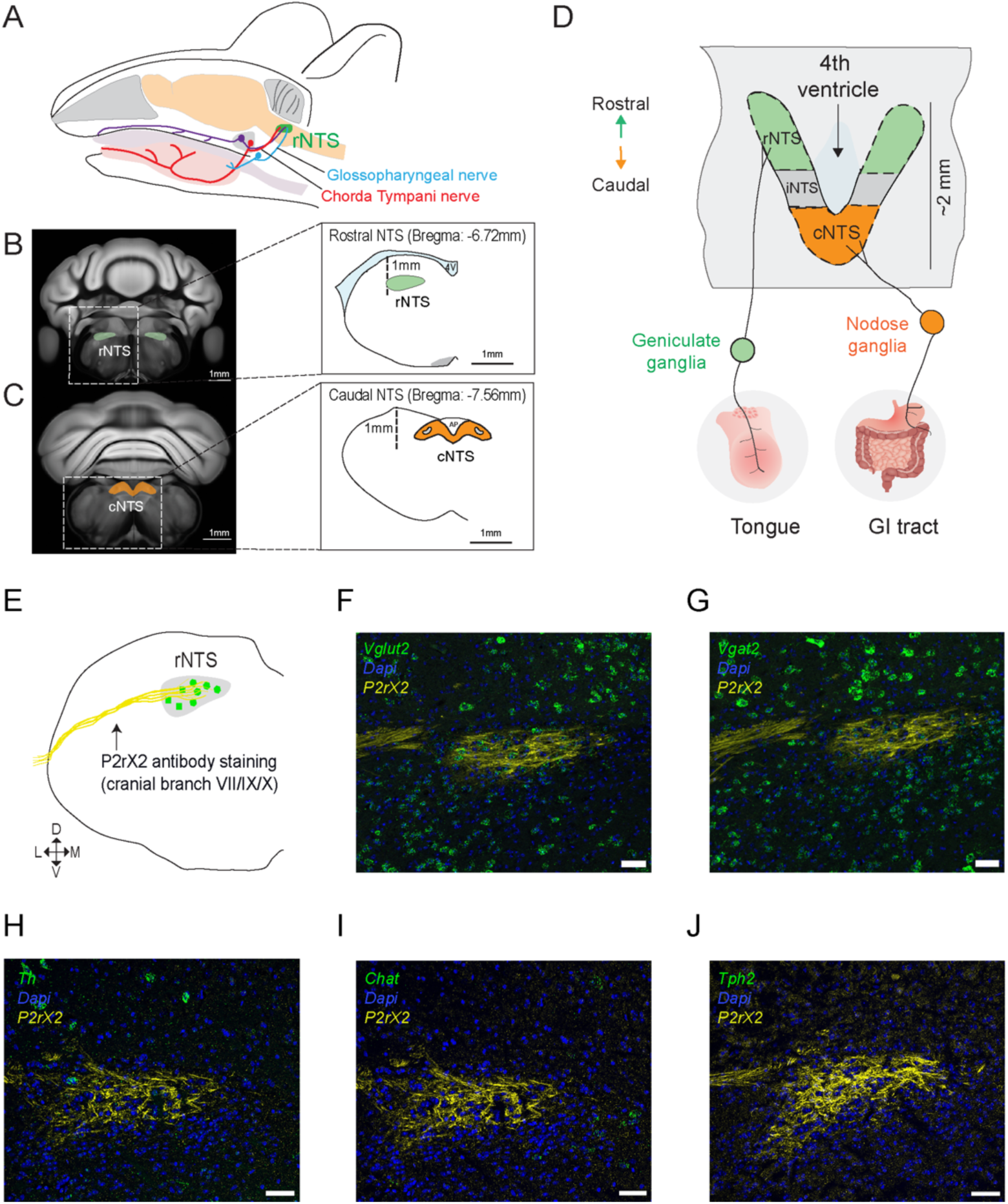
Anatomical and molecular organization of the rostral nucleus of the solitary tract (rNTS). (A) Schematic showing the location of the rNTS within the mouse brainstem and the sensory afferents it receives. (B) Coronal section of the adult mouse brain highlighting the location of the rNTS in the brainstem (left), with corresponding stereotaxic coordinates (right). (C) Coronal section highlighting the location of the caudal NTS (cNTS) in the brainstem (left), with corresponding stereotaxic coordinates (right). (D) Horizontal schematic of NTS subregions and their afferent inputs from the geniculate and nodose ganglia. Molecular organization of the rostral nucleus tractus solitarius (rNTS). Multiplexed RNAscope *in situ* hybridization mapping the principal neurochemical cell types of the rNTS relative to E) the P2rX2-immunolabeled boundaries of the rNTS. RNAscope probes for F) *Slc17a6 (glutamatergic),* G) *Slc32a1 (GABAergic),* H) *Tph2 (serotonergic),* I) *Th (*catecholaminergic), and J) *Chat (*cholinergic) neurons, each co-stained with P2rX2 (yellow). Scale bars, 50 µm. n = 2-3 animals, 6-8 sections.

Metabolic state is communicated to the brain via vagal sensory neurons and circulating hormonal and nutrient signals, including leptin, insulin, ghrelin, glucagon-like peptide-1, and cholecystokinin^9,10^. Glucagon-like peptide-1 (Glp-1) acts both as a circulating gut-derived signal and as a central neuropeptide released by pre-proglucagon neurons resident within the NTS itself^11–13^. Many of these modulators act on receptors within the NTS, most prominently the cNTS, where they regulate meal size, food intake, and long-term energy balance^14,15^. Peripheral taste is similarly state-dependent: leptin acts through Ob-Rb and K-ATP channels in T1R3^+^ taste cells to selectively suppress responses to sweet, but not bitter or sour, compounds^16^, and fasting upregulates glucokinase expression in circumvallate taste bud cells, where it contributes to glucose appetite^17^. State-dependent changes are also detectable centrally: taste responses of rNTS neurons are blunted in diet-induced obese animals compared to lean controls^18^. Together, these findings establish that gustatory signals are modulated by metabolic state at multiple levels, but they leave open the question of where that modulation is imposed. Central changes could be passively inherited from an already modulated periphery, or the rNTS could itself be an independent site of state-dependent regulation. Whether rNTS neurons are themselves equipped to sense metabolic state, and whether their molecular properties are altered when that state changes, remain poorly understood.

Here, we optimized cell-type-specific nuclear RNA sequencing from the rNTS to investigate whether food deprivation alters the transcriptomes of excitatory and inhibitory rNTS neurons. Using Cre-dependent Sun1-sfGFP nuclear tagging coupled with fluorescence-activated nuclear sorting (FANS), we profiled genetically defined excitatory (Vglut2^+^) and inhibitory (Vgat^+^) neurons from ad libitum–fed and 24-hour–fasted mice. Our results show that Vglut2^+^ and Vgat^+^ rNTS neurons are transcriptionally distinct at baseline, expressing largely non-overlapping complements of transcription factors and neuropeptides, and differing in their complement of metabolic-sensing machinery. Superimposed on this baseline divergence, we found that 24-hour fasting elicits a transcriptional response largely restricted to the Vglut2^+^ population, whereas the Vgat^+^ population is minimally affected. Among the genes downregulated in fasted Vglut2^+^ neurons is *Gabrd*, which encodes the δ subunit of the GABA_A_ receptor and is a defining component of the extrasynaptic receptors that mediate tonic inhibition^19^. Together, these findings identify the excitatory rNTS population as the principal transcriptional target of energy deficit and point to the downregulation of tonic inhibitory tone as a cell-type-specific molecular mechanism by which internal state could gate sensory processing at the first gustatory relay in the brainstem.

## Results

### Anatomical and molecular organization of the rNTS neurons

To define the neurochemical composition of the rNTS, we performed multiplexed RNAscope in situ hybridization for markers of the five major neurotransmitter classes present in the brainstem: glutamatergic (*Slc17a6/Vglut2*), GABAergic (*Slc32a1/Vgat*), serotonergic (*Tph2*), catecholaminergic (*Th*), and cholinergic (*Chat*). Because the rNTS lacks clear cytoarchitectonic borders and is contiguous with several functionally distinct nuclei, we combined this panel with immunolabeling for the purinergic receptor P2X2 (P2rX2)^3,20^, which is enriched in primary gustatory and vagal afferent terminals and therefore delineates the incoming afferent field and the boundaries of the solitary tract (Figure 1E). P2rX2 immunoreactivity provided an anatomical fiducial in every section, allowing us to restrict quantification to the region receiving primary gustatory input rather than to a coordinate-defined volume, and to exclude neighboring structures that would otherwise contaminate a purely stereotaxic sampling scheme. Within this P2rX2-defined field, Vglut2^+^ and Vgat^+^ neurons were extensively intermingled throughout the rostrocaudal and mediolateral extent of the rNTS, rather than segregating into discrete subnuclei or laminae (Figure 1F-1G). In contrast, catecholaminergic (*Th*), cholinergic (*Chat*), and serotonergic (*Tph2*) markers were sparse to absent within the rNTS (Figure 1H-1J). The scarcity of Th^+^ neurons is consistent with the known caudal restriction of the A2 noradrenergic group, which lies posterior to the gustatory rNTS, and Chat signal in our sections was confined to the adjacent dorsal motor nucleus of the vagus, providing an internal positive control confirming that the absence of Chat labeling within the rNTS reflects a genuine lack of cholinergic neurons rather than a failure of probe performance. Together, these results establish that excitatory (Vglut2^+^) and inhibitory (Vgat^+^) neurons constitute the two principal cell classes of the rNTS as previously shown by immunohistochemistry^20^. Because these neurons are spatially interleaved, they cannot be separated by microdissection, and bulk profiling of the rNTS necessarily averages the two populations. We therefore turned to a genetic strategy to independently assess the transcriptional profile of each cell class.

### Cell-type-specific nuclear RNA-sequencing of excitatory and inhibitory neurons in rNTS

We then designed a nuclear RNA-sequencing workflow to profile excitatory (Vglut2^+^) and inhibitory (Vgat^+^) populations across metabolic states (Figure 2A). To gain genetic access to each population, we crossed *Vglut2-Cre* and *Vgat-Cre* driver lines to the Cre-dependent Sun1-sfGFP reporter, generating Vglut2-Cre/+; Sun1-sfGFP/+ and Vgat-Cre/+; Sun1-sfGFP/+ mice in which the nuclear envelope of excitatory and inhibitory neurons, respectively, is marked with GFP (Figure 2B-2C). Because the Sun1 fusion protein is retained at the inner nuclear membrane, GFP labeling survives nuclear isolation, allowing genetically defined nuclei to be recovered from tissue where the two populations are anatomically inseparable^23,24^. rNTS tissue was microdissected from ad libitum-fed and 24-hour-fasted mice of both genotypes, nuclei were isolated under RNase-free conditions, and Sun1-sfGFP^+^ nuclei were purified by fluorescence-activated nuclei sorting (FANS) using a sequential gating strategy: intact singlet nuclei were first selected on the basis of forward and side scatter, followed by selection of the GFP^+^ fraction (Figure 2D–2E). Sequencing yielded a mean of 34.6 million reads per library after adapter and quality trimming (*Vglut2^+^*, 41.9M; *Vgat^+^*, 26.1M), the large majority of which passed filter and aligned uniquely to the mouse genome (STAR; mean 79.8% uniquely mapped across all 13 libraries, range 66.4–89.3%). Libraries from both genotypes and both feeding conditions showed comparable mapping rates, duplication rates (∼30-45%), and gene-body coverage (∼70- 80%) across both genotypes, indicating that downstream differences between groups are unlikely to reflect systematic variation in library quality (Figure S1). Excitatory and inhibitory rNTS neurons showed markedly different global transcriptional profiles. Principal component analysis of variance-stabilized expression resolved the two populations along the first principal component, which alone captured 81.9% of the variance and separated Vglut2^+^ from Vgat^+^ nuclei into non-overlapping regions, while the second component (7.8%) reflected finer within-population structure (Figure 2F). This separation exceeded the effect of feeding condition: fed and fasted samples of the same cell type did not resolve along the first two components, confirming that the sort captured two genuinely distinct populations. We next compared the transcriptional response to 24 hour fasting between excitatory and inhibitory rNTS populations. Across the whole tested transcriptome, the distribution of fasting log₂ fold changes differed significantly between the two populations, with the Vglut2^+^ distribution carrying a heavier tail of responsive genes (Figure 2G) (two-sample Kolmogorov-Smirnov D = 0.351; because genes are not independent observations, we report this as a descriptive comparison of the two response distributions rather than a formal test; nominal p < 2.2 × 10⁻¹⁶). Genotype specificity of the sorted material was confirmed at the transcript level: Vglut2^+^ was strongly enriched in excitatory neuron libraries and Vgat^+^ in inhibitory neuron libraries (Figure 2H), establishing that FANS recovered the intended populations. When genes were grouped into functional classes, the two populations differed in their baseline complement of metabolic machinery: insulin signaling and lipid metabolism genes showed a significant baseline bias between cell types, whereas hunger-sensing, satiety-sensing, and energy-sensing classes did not (Figure 2H). Excitatory and inhibitory rNTS neurons are therefore not merely transcriptionally distinct in general; they are differently equipped to read the metabolic state of the animal.

**Figure 2.**
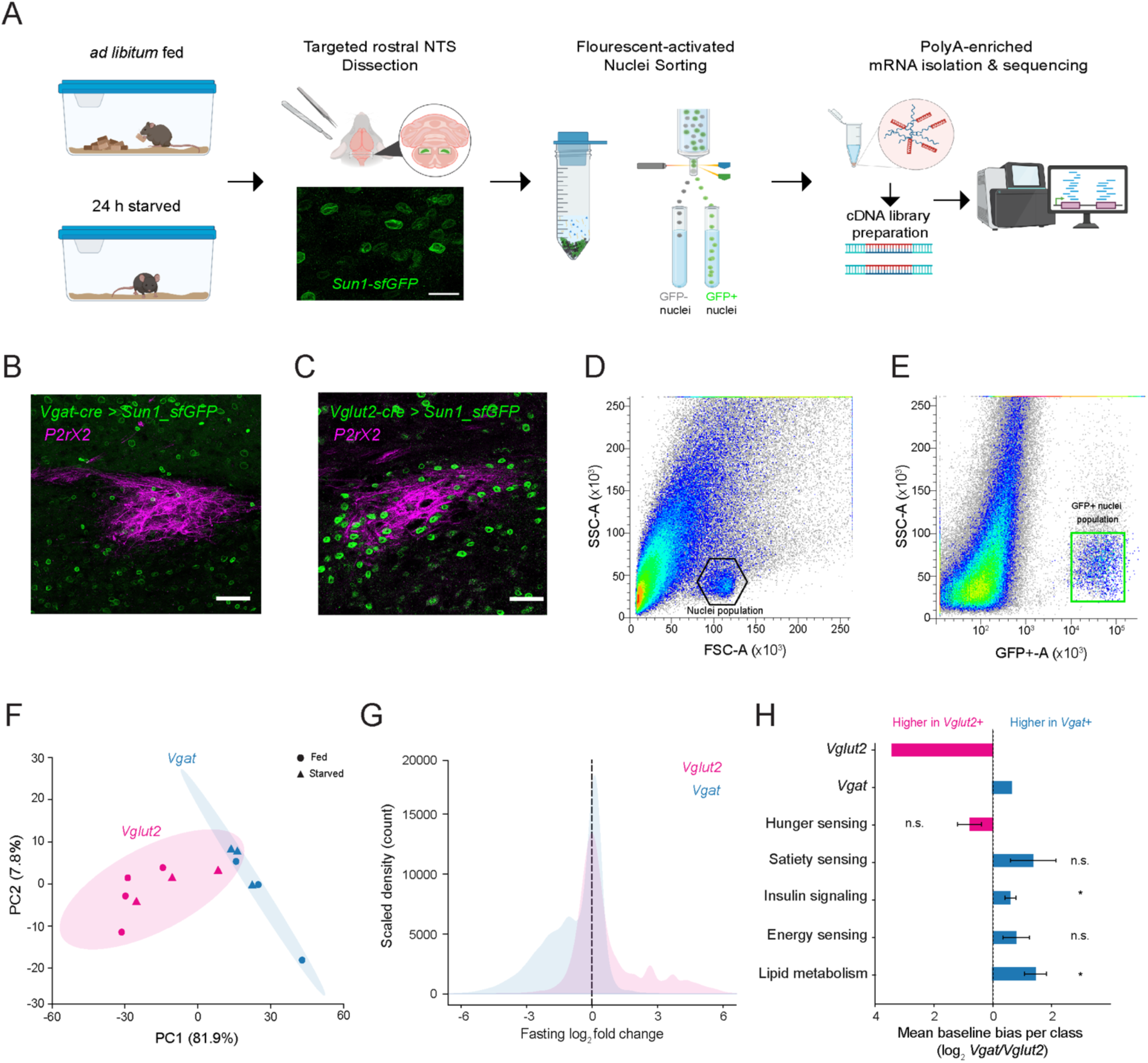
Nuclear RNA-sequencing of genetically defined rNTS neurons. (A) Experimental workflow: ad libitum-fed or 24-h-fasted *Vglut2*^Cre-/+^; Sun1-sfGFP and *Vgat* ^Cre-/+^; Sun1-sfGFP mice underwent targeted rNTS microdissection (representative image of Sun1-sfGFP nuclei, scale bar, 20 um), nuclei isolation, fluorescence-activated nuclei sorting (FANS) of Sun1-sfGFP^+^ nuclei, polyA-enriched mRNA isolation, cDNA library preparation, and RNA-sequencing. (B, C) Representative confocal images of Sun1–sfGFP nuclear-envelope labeling (green) in the rNTS of *Vglut2*^Cre-/+^; Sun1-sfGFP (*Vglut2-cre>Sun1-GFP*) (B) and *Vgat* ^Cre-/+^; Sun1-sfGFP (*Vgat-cre>Sun1-GFP*) (C) mice, co-stained for P2rX2 (magenta). Scale bars, 50 µm. (D) FANS gating for intact nuclei by forward/side scatter. (E) GFP^+^ nuclei gated for collection using the BD FACSMelody™ cell Sorter purity mode) with a 100 µm nozzle chip. (F) PCA of variance-stabilized expression; each point is one library, colored by cell type (*Vglut2* - pink, *Vgat* - blue) and shaped by condition (circle, fed; triangle, fasted); shaded ellipses, 95% confidence regions; PC1, 81.9%; PC2, 7.8%. (G) Scaled-density (count) distributions of fasting log₂ fold change (fasted/fed) across all tested genes, compared by two-sample Kolmogorov - Smirnov test (D = 0.351, p < 2.2 × 10^-16^). (H) Mean baseline (fed) cell-type bias per functional class, log₂(*Vgat/Vglut2*); bars, class mean (right/blue, higher in *Vgat*; left/pink, higher in *Vglut2*); points, individual genes; error bars, SEM; identity markers *Vglut2* and *Vgat* shown separately; asterisks indicate significant differences (binomial sign test, n.s., not significant)

### Excitatory and inhibitory rNTS neurons differ in their metabolic-sensing machinery

To examine the transcriptional differences between Vglut2^+^ and Vgat^+^ neurons at the level of individual genes, we compared the expression of hormone receptors, neuropeptides, and nutrient-sensing effectors across the two populations. Control genes behaved as expected: *Gapdh* was robustly expressed in both populations, and Vglut2 and Vgat transcripts were each enriched in the corresponding genotype (Figure 3A), consistent with recovery of the intended populations by FANS. We next surveyed the major classes of metabolic genes in each population. Both cell types expressed a broad complement of metabolic hormone receptors, including *the leptin receptor (Lepr)*, *the insulin receptor (Insr)* together with its downstream substrates (*Irs1, Irs2*), t*he IGF-1 receptor (Igf1r)*, *the GLP-1 receptor (Glp1r)*, and both *adiponectin receptors (Adipor1, Adipor2)* (Figure 3B). The rNTS is therefore positioned to read out multiple circulating signals of energy availability, and to do so through both of its principal cell classes. The Vglut2^+^ and Vgat^+^ populations were not equivalently equipped, however. *Igf1r* and *Insr* were expressed at substantially higher levels in Vgat^+^ than in Vglut2^+^ neurons, consistent with the significant baseline bias of the insulin-signaling gene class toward the inhibitory population (Figure 2H). *Cckbr* was detected only in Vglut2^+^ neurons, whereas *Glp2r* was detected only in Vgat^+^ neurons. Two receptors were conspicuously absent. Neither *the ghrelin receptor (Ghsr)* nor *the GIP receptor (Gipr)* was detected above threshold in either population, indicating that the rNTS is not a direct target of ghrelin or of gastric inhibitory polypeptide, and that any influence of these hormones on gustatory processing must be relayed indirectly, for example, through the vagal afferents or the cNTS. The asymmetry was more pronounced among neuropeptide-related genes (Figure 3C). *Cck, Mc4r, Mchr1*, and *Npy1r* were all detected in Vglut2^+^ neurons but not in Vgat^+^ neurons, indicating that the neuropeptidergic arm of metabolic signaling converges preferentially on the excitatory population. Pomc was absent from both populations, consistent with the rNTS lacking a resident population of POMC neurons.

**Figure 3.**
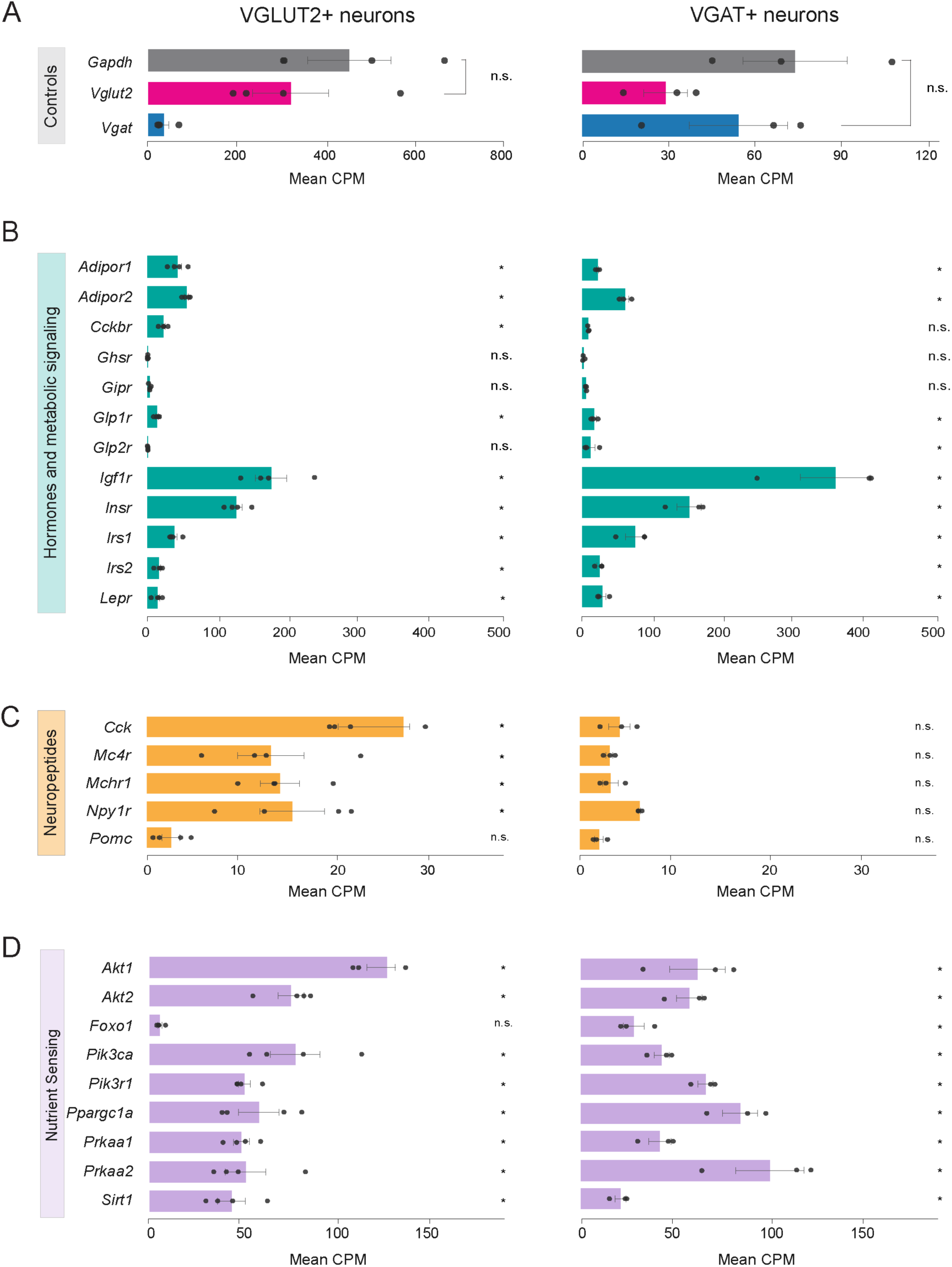
Baseline expression of identity, hormone/metabolic, neuropeptide, and nutrient-sensing genes. (A-C) Mean expression (CPM) under the ad libitum-fed condition for VGLUT2^+^ (left) and VGAT^+^ (right) populations; points, individual replicates; error bars, SEM; asterisks denote genes above a 10 CPM detection threshold; n.s., not significant. (A) Control/identity genes (*Gapdh, Vglut2, Vgat).* (B) Hormone and metabolic-signaling receptors. (C) Neuropeptides and neuropeptide receptors. (D) Nutrient- and energy-sensing genes. Note the independent x-axis scale per row and panel. n = 3-4 fed per population.

In contrast to hormone and peptide signaling pathways, the intracellular nutrient-sensing machinery was broadly and comparably expressed in both cell types (Figure 3D). Components of the insulin–PI3K–AKT cascade (*Pik3ca, Pik3r1, Akt1, Akt2, Foxo1*) and of the AMPK and sirtuin pathways (*Prkaa1, Prkaa2, Sirt1, Ppargc1a*) were present in Vglut2^+^ and Vgat^+^ neurons alike, indicating that both populations retain the capacity to transduce a metabolic signal once it is received. The divergence between the two classes therefore lies not in the intracellular machinery for interpreting energy status, but in the receptors that determine which extracellular signals reach them in the first place. Taken together, these data demonstrate that both rNTS cell classes are equipped to sense circulating metabolic signals, but that they are differently tuned to the signals they receive: hormonal input is biased toward inhibitory neurons, and neuropeptidergic input toward excitatory neurons.

### Fasting minimally alters the transcriptional profile of inhibitory rNTS neurons

We next compared the transcriptional response to 24-hour fasting between excitatory and inhibitory rNTS populations. We first asked how Vgat^+^ neurons respond to 24 hours of food deprivation. Housekeeping genes (*Gapdh, Actb*) and the cell-type marker *Slc32a1* were unchanged between fed and fasted animals, confirming that the libraries were comparable across conditions (Figure 4A). Fasting produced a minimal transcriptional response: identity and housekeeping genes were stable across conditions (Figure 4A). Differential expression analysis revealed a strikingly muted response. Only a single gene, *Atp11b*, reached significance at FDR < 0.10, with three further genes (*Ipo9, Ddx17, Phka4*) passing an exploratory threshold of FDR < 0.25 (Figure 4B). Because a transcriptional response could in principle be functionally meaningful even if it involved few genes, we examined the core GABAergic program directly. Neither the GABA synthesis and transport machinery (*Gad1, Gad2, Slc32a1, Slc6a1, Slc6a11, Slc6a12, Slc6a13, Aldh5a1, Abat*) (Figure 4C) nor the presynaptic release machinery (S*yt1, Syt2, Snap25, Vamp2, Stx1a, Stx1b, Stxbp1, Cplx1, Cplx2, Rims1, Rims2, Rab3a, Unc13a*) (Figure 4D) showed any fasting-induced change. The full complement of GABA_A_ receptor subunits expressed in Vgat^+^ neurons was likewise unaffected (Figure 4E - 4F). Thus, at the transcriptional level, inhibitory rNTS neurons appear to be largely indifferent to a 24-hour energy deficit.

**Figure 4.**
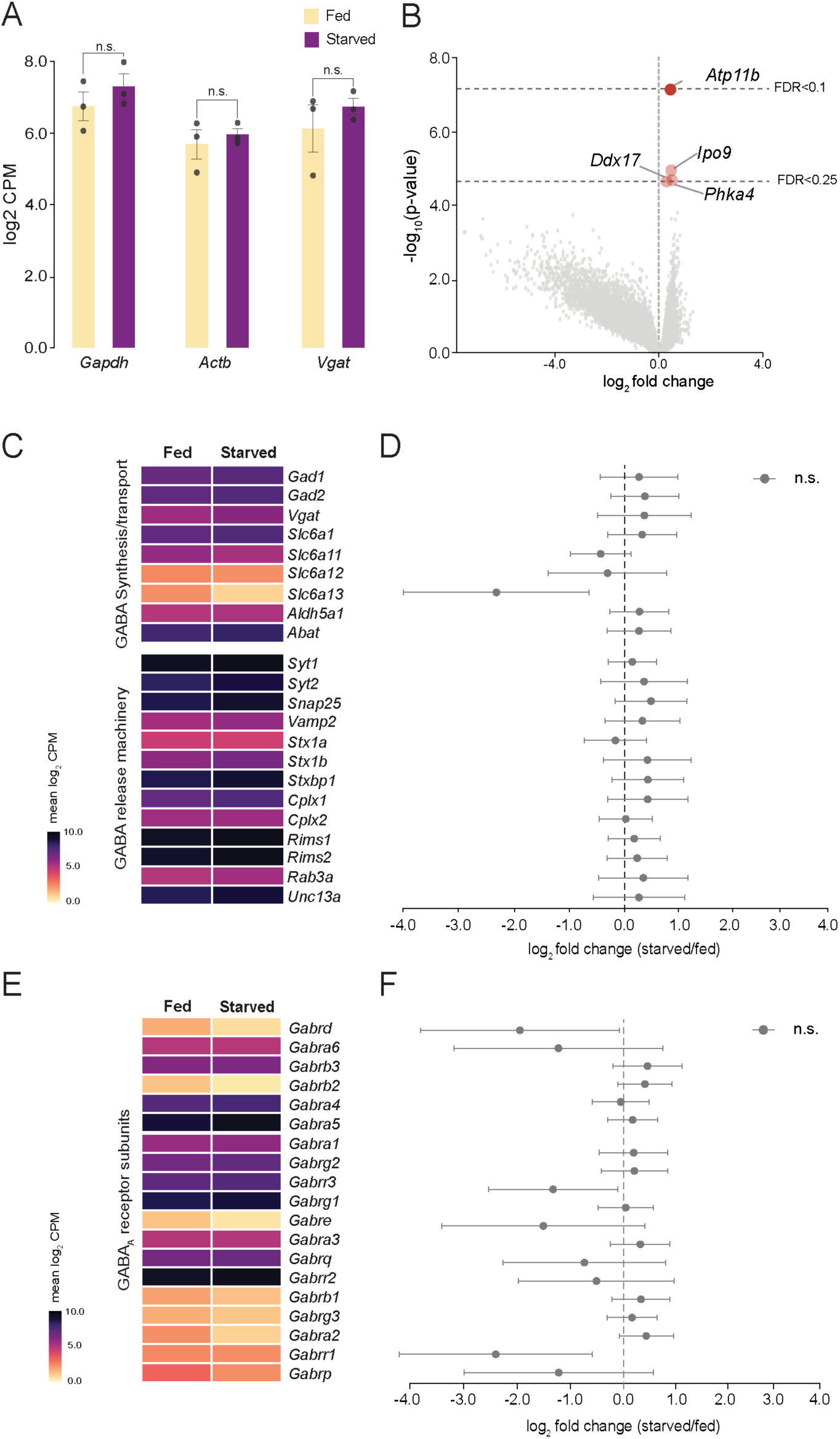
The inhibitory (Vgat^+^) population is transcriptionally unresponsive to fasting. (A) Housekeeping/identity gene expression (log₂ CPM; *Gapdh, Actb, Vgat)* in fed (yellow) vs fasted (deep purple); bars, mean; points, replicates; error bars, SEM; all n.s. (B) Volcano plot of the Vgat^+^ neuronal fasting response; dashed lines mark nominal p corresponding to FDR = 0.10 and 0.25; *Atp11b* is the only gene at FDR < 0.10; genome-wide BH-adjusted p-values using DESeq2 analysis. (C-D) GABA synthesis/transport and vesicular release machinery: heatmap of mean log₂ CPM (fed, fasted) with forest plot of fasting log₂ fold change (point, MLE; whiskers, 95% CI; blue, FDR < 0.10; grey, n.s.). (E-F) GABA-A receptor subunits, as in C; no subunit reached FDR < 0.10. n = 3 ad libitum fed; 3 24 h fasted.

### Fasting selectively downregulates *Gabrd* expression in excitatory rNTS neurons

We next performed the same analysis in Vglut2^+^ neurons. As in Vgat^+^ neurons, the housekeeping genes *Gapdh* and *Actb* and the cell-type marker *Slc17a6* were stable across feeding conditions, indicating that the fed and fasted libraries were comparable (Figure 5A). Differential expression analysis, however, revealed a substantially larger response. Two genes reached significance at FDR < 0.10, *Gabrd* and *Uqcrc2*, and approximately twenty further genes passed the exploratory threshold of FDR < 0.25, including *Fh1*, *Atp5pb*, *Rraga*, *Ier5*, and several chaperonin subunits (*Cct3*, *Cct5*, *Tcp1*) (Figure 5B). Both *Gabrd* and *Uqcrc2* were downregulated in the fasted state. Several of these genes are functionally related. *Uqcrc2* encodes a core subunit of mitochondrial complex III*; Fh1* encodes fumarate hydratase; *Atp5pb* encodes a subunit of ATP synthase; and *Rraga* encodes a Rag GTPase that couples amino acid availability to mTORC1^21^. Their appearance among the fasting-responsive genes suggests a coordinated adjustment of oxidative metabolism and nutrient sensing in excitatory rNTS neurons and is consistent with a cell-autonomous response to systemic energy deficit. We note that this signature was absent from the Vgat^+^ population, which showed no metabolic genes among its few differentially expressed transcripts.

**Figure 5.**
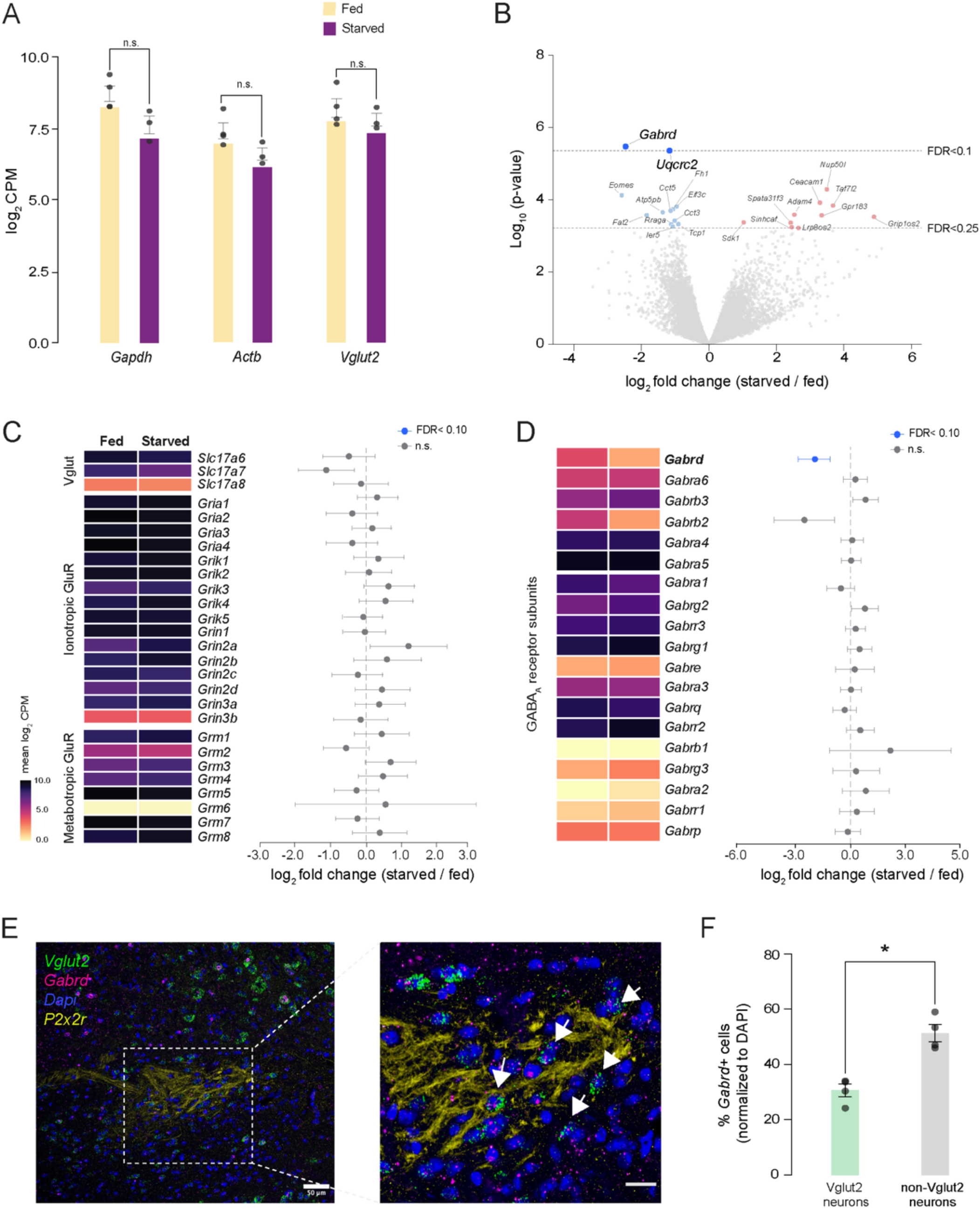
Fasting selectively reduces Gabrd-mediated tonic-inhibition machinery in excitatory (Vglut2^+^) neurons. (A) Housekeeping/identity gene expression (log₂ CPM; *Gapdh, Actb, Vglut2)* in fed (yellow) vs fasted (deep purple); all n.s. (B) Volcano plot of Deseq2 differentially expressed genes in fasted vs. fed mice; dashed lines mark FDR = 0.10 and 0.25; *Gabrd* and *Uqcrc2* are the two genes at FDR < 0.10, both downregulated; genome-wide BH-adjusted p-values using DESeq2 analysis. (C) Glutamatergic pathway genes (VGLUT transporters, ionotropic and metabotropic glutamate receptors): heatmap with forest plot of fasting log₂ fold change; none reach FDR < 0.10. (D) GABA-A receptor subunits: heatmap with forest plot of fasting log_2_ fold change; *Gabrd* is the only subunit at FDR < 0.10. (E) Representative RNAscope image of the rNTS (DAPI, blue; *Vglut2,* green; *Gabrd,* magenta; P2rX2, yellow, scale bar 50 um); arrows mark *Vglut2^+^/Gabrd*^+^ cells in the zoomed-in image; scale bar, 20 µm. (F) Percentage of *Gabrd^+^* cells (normalized to DAPI) among *Vglut2^+^* (31.5%) versus non-Vglut2 (51.9%) neurons; bars, mean; points, individual animals; error bars, SEM; *p < 0.05, paired t-test; n = 3 mice; 4 sections.

We further investigated whether glutamatergic signaling is altered in Vglut2^+^ neurons. Neither the vesicular glutamate transporters (*Slc17a6*, *Slc17a7*, *Slc17a8*) nor any AMPA (*Gria1–4*), kainate (*Grik1–5*), NMDA (*Grin1*, *Grin2a–d*, *Grin3a–b*), or metabotropic (*Grm1–8*) glutamate receptor subunit changed their expression levels with fasting (Figure 5C). The machinery of excitatory neurotransmission in these neurons is thus transcriptionally stable across metabolic states, and fasting does not act by adjusting the receptors through which excitatory input is received. We next focused on GABAergic signaling in this population. We found that *Gabrd* was the only GABA_A_ receptor subunit among the nineteen we examined that was significantly altered by food deprivation, getting strongly downregulated upon fasting. Notably, neither Gabra4 nor Gabra6 changed in response to fasting (Figure 5D). Because δ does not form functional receptors on its own, and instead partners obligatorily with α4 or α6, one might expect the α subunits to be co-regulated with it. Our results indicate that the fasting response does not reflect a coordinated transcriptional program acting on the extrasynaptic receptor as a unit, but rather a change in the availability of a single component. This is consistent with the prevailing view that δ, rather than its α partners, is limiting for extrasynaptic receptor assembly: α4 subunits are expressed in excess and can also participate in α4βγ2 synaptic receptors, whereas δ is incorporated only into extrasynaptic complexes. Under this interpretation, a reduction in *Gabrd* alone would be sufficient to lower the number of functional δ-containing receptors at the membrane, and therefore the magnitude of the tonic conductance they carry, without any accompanying change in α subunit expression.

Because our RNA sequencing experiments were performed on nuclei sorted from dissociated tissue, we next confirmed that *Gabrd* is expressed in excitatory rNTS neurons in situ. We performed RNAscope for *Gabrd* together with *Vglut2* within the P2rX2-defined rNTS (Figure 5E). *Gabrd* transcripts were readily detected in *Slc17a6*^+^ neurons, and approximately 31% of Vglut2^+^ cells were *Gabrd*^+^ (Figure 5F). *Gabrd* was also expressed in the non-Vglut2^+^ population, and at a higher frequency (approximately 51%; SEM; *p < 0.05, paired t-test; n = 3 mice; 4 sections), indicating that the subunit is not restricted to excitatory neurons. Together, our results confirm that *Gabrd* is expressed in a substantial fraction of Vglut2^+^ rNTS neurons, and that the downregulation we detected by sequencing is unlikely to reflect contamination of the sorted material. They also make the effect’s selectivity more informative than it would otherwise be. *Gabrd* is expressed in both cell classes and, more broadly, in the non-Vglut2^+^ population, yet only the Vglut2^+^ population downregulates it in response to fasting. The cell-type specificity of the effect therefore does not follow from a cell-type-specific distribution of the transcript. It reflects a difference in how these two rNTS populations regulate a gene they both express.

## Discussion

Recent single-nucleus atlases have mapped the cellular composition of the dorsal vagal complex, identifying dozens of neuronal subtypes across the NTS, area postrema, and dorsal motor nucleus of the vagus^9,10^. These datasets are dominated by the caudal, viscerosensory NTS, are organized around unsupervised clustering rather than around the excitatory-inhibitory division that structures local processing in the gustatory relay, and profile animals in a single metabolic state. Our data are complementary: we sample the rostral, gustatory division specifically, using P2rX2 immunolabeling to restrict dissection to the primary afferent field, and we ask not what cell types exist but how a defined pair of them responds to energy deficit. To our knowledge, the transcriptional response of genetically defined excitatory and inhibitory rNTS neurons to fasting has not previously been described.

In this study, we asked whether metabolic state alters the transcriptional profile of rNTS neurons, and whether any such response is uniform across the nucleus or restricted to particular cell types. Our data provide the first cell-type-resolved molecular description of the gustatory rNTS. We show that the two principal classes of this nucleus, excitatory (Vglut2^+^) and inhibitory (Vgat^+^), are intermingled, similarly positioned within the afferent field. Furthermore, we found that both neuronal populations are equipped to sense circulating metabolic signals and respond to hunger in fundamentally different ways. Using bulk nuclear RNA sequencing, we found that 24 hours of food deprivation reshapes the Vglut2^+^ transcriptome while leaving the Vgat^+^ transcriptome nearly untouched. In particular, fasting selectively affects the expression of *Gabrd* in Vglut2^+^ neurons, which encodes the δ subunit of the GABA_A_ receptor, responsible for extrasynaptic tonic inhibition. Such a targeted, directional response identifies downregulation of the tonic inhibitory machinery as a candidate molecular substrate through which the neurophysiology of excitatory rNTS neurons may be affected.

GABA_A_ receptors are persistently activated by the low concentrations of ambient GABA in the extracellular space and generate a tonic chloride conductance, rather than the fast phasic currents produced by synaptic receptors. Because tonic inhibition governs neurophysiological gain control rather than discrete, momentary synaptic events, it is a strong candidate mechanism for modulating sensory responses in both cortical and subcortical regions of taste processing^22–25^. δ-GABA_A_ receptors have recently been shown to gate sweet preference in the adult mouse gustatory insular cortex, where their neurosteroid sensitivity confers state-dependent control over ingestive choice^26^. Our data place an analogous mechanism at the opposite end of the taste axis: the same subunit in the first central relay is under transcriptional control by metabolic state. That δ-containing receptors modulate taste-guided behavior at both the earliest and the latest stages of central processing suggests that tonic inhibition may be a recurring motif for state-dependent gain control throughout the gustatory system. Functional studies in the rNTS support this interpretation. Enhancing GABAergic tone in the rNTS of awake, licking rats reconfigures sensorimotor activity and remodels the population representation of taste, including its dynamic encoding of palatability^26^. Conversely, blocking GABA_A_ receptors in the NTS broadens the tuning of taste-responsive neurons and increases the heterogeneity of their responses^24^. Together, these pharmacological manipulations establish that the level of GABA_A_ receptor-mediated inhibition is not a passive background constraint but an active determinant of how rNTS circuits represent taste identity and hedonic value, and that shifting it in either direction is sufficient to reshape that representation. What these experiments could not address is how such a shift might be produced endogenously. Both manipulations were acute and pharmacological; both acted indiscriminately on synaptic and extrasynaptic receptors; and neither could distinguish the cell type in which inhibition was altered. Our data speak to each of these points. We find that fasting selectively reduces *Gabrd*, the subunit that confers extrasynaptic localization and mediates tonic rather than phasic inhibition, and that it does so specifically within the excitatory population that carries gustatory information forward from the first central relay. The remaining GABA_A_ subunits, including those that assemble into the canonical synaptic receptor, are unchanged, as are all glutamate receptor subunits.

Together, our data identify a candidate molecular, cell-type-specific substrate through which the animal’s own metabolic state could impose the kind of shift in inhibitory tone that these pharmacological studies induced exogenously. In a fasted animal, a reduction in the tonic conductance carried by δ-containing receptors would be expected to lower the gain of excitatory rNTS neurons’ input-output function, increasing their responsiveness to a given amount of afferent drive. The predicted consequence of this regulation would be enhanced central taste responses under energy deficit, which is the direction required to account for the increased behavioral salience of palatable tastants during hunger, and it places the mechanism at the earliest stage at which central gain control is possible.

## Acknowledgements

We thank members of the Yapici and Markenscoff-Papadimitriou laboratories for discussions and comments on the manuscript. We are grateful to Jen Grenier for advice on nuclear isolation and fluorescence-activated nuclei sorting, and to the Cornell Institute of Biotechnology Transcriptional Regulation and Expression (TREx) Facility for library preparation and sequencing. We thank the Cornell Institute of Biotechnology Flow Cytometry Facility for assistance with nuclei sorting, and the Cornell Center for Animal Resources and Education for animal care. Figure 2A was created in part with BioRender. This work was supported by a seed grant from the Cornell Center for Vertebrate Genomics to N.Y. and E.M.-P.; a Brain Research Foundation Seed Grant and NIH grant 1R34NS144924 to N.Y.; and a Simons Foundation grant SFARI 868055 to E.M.-P. N.Y. is a Nancy and Peter Meinig Family Investigator in the Life Sciences.

## Author Contributions

D.M., E.M.P., and N.Y. conceived the study and designed the experiments. D.M. performed all experiments, including the RNAscope in situ hybridization, rNTS microdissection, nuclear isolation, fluorescence-activated nuclei sorting, and library preparation. D.M. and E.M.-P. performed the RNA-sequencing analysis. D.M. and N.Y prepared the figures. D.M. wrote the manuscript with input from N.Y. and E.M.P. N.Y. and E.M.-P. supervised the project and acquired funding. All authors read and approved the final manuscript.

## Competing Interests

The authors declare no competing interests.

## Declaration of Generative AI Use

During the preparation of this work, the authors used Claude (Anthropic) to edit code and improve the language, clarity, and readability of the manuscript. After using these tools, the authors reviewed and edited the content as needed and took full responsibility for the content.

---

**Supplementary Figure S1.**
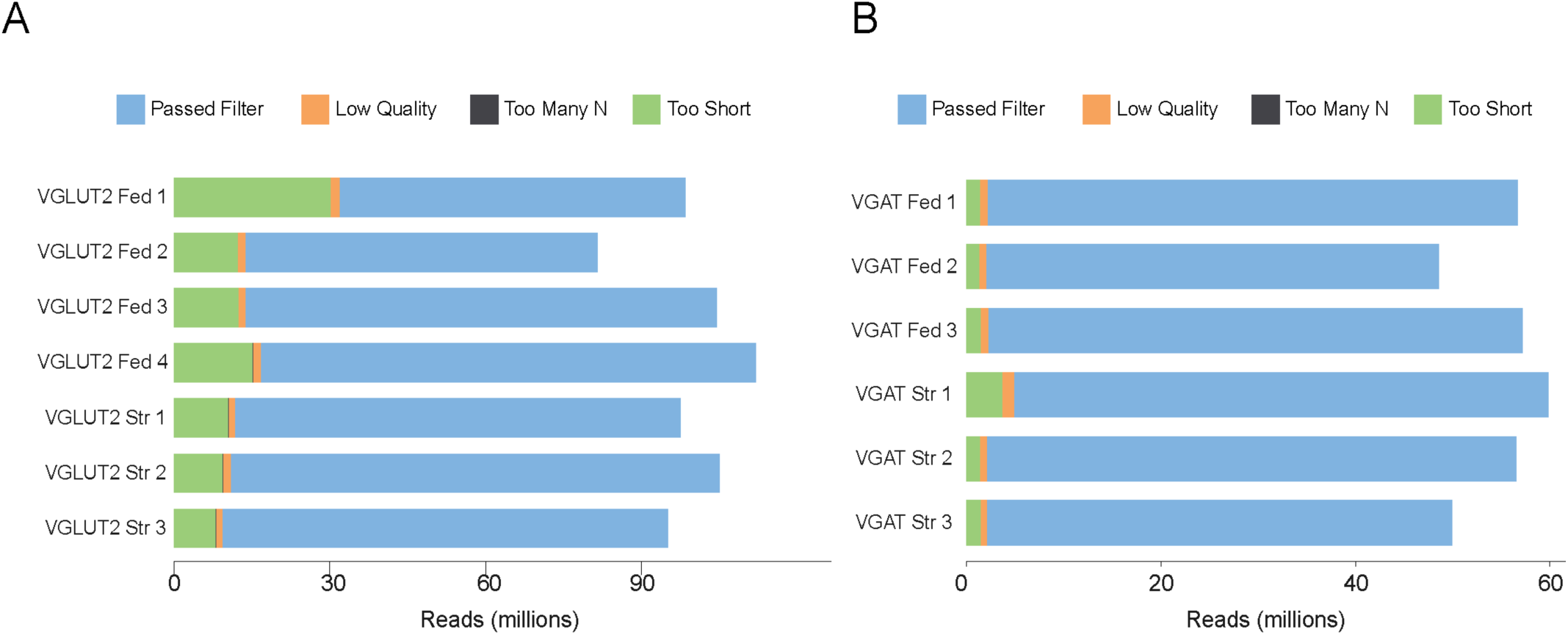
Sequencing read quality control. Read filtering summary (fastp) for each sequencing library, showing the number of reads passing filter and those removed for low quality, excessive ambiguous bases (too many N), or insufficient length (too short), in millions of reads. (A) Vglut^+^ libraries (4 *ad libitum*–fed, 3 fasted). (B) Vgat^+^ libraries (3 *ad libitum*–fed, 3 fasted). Most reads passed filter across all libraries.

## Materials and Methods

### Animals

All procedures were approved by the Institutional Animal Care and Use Committee of Cornell University and performed in accordance with the NIH Guide for the Care and Use of Laboratory Animals. RNAscope in situ hybridization experiments were performed in wild-type *C57BL/6J* mice (JAX #000664). For cell-type-specific nuclear isolation, homozygous *Vglut2*^Cre/Cre^(*Slc17a6^tm2(cre)Lowl^*/J) JAX#016963 or *Vgat*^Cre/Cre^ (*Slc32a1^tm2(cre)Lowl^*/J) JAX#016962 mice were crossed to the Cre-dependent CAG-Sun1-sfGFP reporter (*ROSA26*^CAG-Sun1–sfGFP–Myc^) (JAX #021039), in which a loxP-flanked stop cassette prevents expression of a Sun1–sfGFP–Myc fusion protein until Cre-mediated recombination. Cre-dependent excision drives expression of the Sun1–sfGFP fusion at the inner nuclear membrane, labeling the nuclear envelope of *Vglut2^+^* (excitatory) or *Vgat^+^* (inhibitory) neurons. Experimental animals were hemizygous for both alleles. Genotypes were confirmed by PCR using JAX-recommended primers. Male and female mice aged 8–12 weeks were used in equal proportions across all experiments. Animals were group-housed on a 12:12 light: dark cycle, with ad libitum access to standard chow and water, except during fasting. Fasted mice underwent 24 hours of complete food deprivation with continued water access; controls remained *ad libitum* fed. Fasting was initiated at ZT14 for *Vglut2*^Cre/+^; *Sun1-sfGFP^1^*^x/+^ and ZT17 for *Vgat*^Cre/+^; *Sun1-sfGFP*^1x/+^ mice on the previous day, and tissue was collected at the same ZT on the day of experiments across conditions to control for circadian effects.

### RNAscope in situ hybridization

Wild-type adult C57BL/6 J mice (4 males and 2 females, 20–25 g) aged 8-14 weeks were used for RNAscope *in situ* hybridization experiments. Mice were deeply anesthetized with Pentobarbital (Fatal Plus, 035946) at a concentration of 100 mg/kg body weight of adult mice and transcardially perfused with PBS followed by 4% paraformaldehyde in 0.1 M PBS. Brains were post-fixed for 12- 16 hours at 4 °C, cryoprotected in 30% sucrose in PBS until equilibrated for up to 48 h, embedded in OCT, and frozen on dry ice. Coronal sections (8 µm) spanning the rostral NTS were collected on a sliding microtome (Leica SM2000R) onto SuperFrost Plus slides and stored at −80 °C. Multiplexed fluorescent in situ hybridization was performed using the RNAscope Multiplex Fluorescent Reagent Kit v2 with TSA Vivid dyes (Advanced Cell Diagnostics, 323100 and 323270) according to the manufacturer’s protocol for fixed-frozen tissue. Probes were: *Slc17a6* (*Mm-Slc17a6-C2, 319171-C2*), *Slc32a1* (*Mm-Slc32a1-C3, 319191-C3*), *Tph2* (*Mm-Tph2-C2, 318691-C2*), *Th* (*Mm-Th-C3, 317621-C3*), *Chat* (*Mm-Chat-C2, 408731-C2*), and *Gabrd* (*Mm-Gabrd-C3, 459481-C3*). Because several probes share a channel, they were assigned to non-overlapping multiplex panels. Slc17a6/Tph2/Chat are all C2 and Slc32a1/Th/Gabrd are all C3. During the experiments, rNTS was delineated by immunolabeling for the P2rX2 receptor, which marks gustatory afferent terminals in the rostral NTS. Following the final HRP-blocking step of the RNAscope protocol, sections were processed for immunofluorescence using the ACD Co-Detection workflow. Sections were blocked and incubated with rabbit anti-P2rX2 (Alomone Labs, APR-003; 1:1000) overnight at 4°C, followed by incubation with goat anti-rabbit Alexa 647 secondary antibody (Thermo Scientific, A-21244; 1:2000) for 3 hours at room temperature. Nuclei were counterstained with DAPI, and sections were coverslipped in Slowfade™ gold antifade mountant (Thermo Scientific, S36936). Images were acquired on a Leica Stellaris 3 confocal microscope utilizing the Leica Application Suite X software with an HC PL APO 20x/0,75 CS2 or HC PL APO 63x/1,40 OIL CS2 objectives at a pixel resolution of 3128 x 2568 px (63x Objective), 5296 x 5296 pixels (20x objective) and 5-6 um z-stacks at a single slice thickness of 0.5 um. Acquisition settings were held constant across conditions within an experiment.

### Nuclei isolation and fluorescence-activated nuclei sorting (FANS)

*Vglut2*^Cre/+^; *Sun1-sfGFP^1^*^x/+^ or *Vgat*^Cre/+^; *Sun1-sfGFP*^1x/+^ mice (4–5 animals pooled per sample) were euthanized, and brains were rapidly extracted. Coronal sections (500 µm) spanning the rostral NTS were cut on ice in an adult mouse brain matrix (Kent Scientific), and the rNTS was bilaterally microdissected under a stereomicroscope using landmarks such as the rostral pole of the area postrema and the fourth ventricle as boundaries. Dissected tissue was collected into 1 ml of ice-cold homogenization buffer I (250 mM sucrose, 25 mM KCl, 5 mM MgCl₂, 120 mM tricine-KOH, pH 7.8) supplemented with 100 µM dithiothreitol, 100 µM phenylmethylsulfonyl fluoride, 0.4 U/ml SUPERase·In RNase inhibitor (Thermo Fisher, AM2694), and 1x EDTA-free protease inhibitor cocktail (Sigma-Aldrich, 11873580001). Tissue was homogenized with 10 strokes of a loose pestle in a 1 ml Dounce glass tissue grinder. An equal volume (1 ml) of homogenization buffer II (homogenization buffer I, 0.4% NP-40) was then added, and the suspension was gently pipetted multiple times and incubated on ice for 10 min. The homogenate was centrifuged at 1,000 × g for 10 min at 4 °C, and the nuclear pellet was resuspended in 0.2 ml of sorting buffer (1x phosphate-buffered saline, fetal bovine serum (A5670701), 0.15 mM spermine, 0.5 mM spermidine, 0.4 U/ml SUPERase.In, and 1× EDTA-free protease inhibitor cocktail). GFP^+^ nuclei (80,000–100,000 per sample) were sorted directly into TRIzol™ LS reagent solution (Thermo Fisher; 10296010) on a BD FACSMelody™ Cell Sorter (purity mode) using a 100 µm nozzle chip. Sorted nuclei in TRIzol were flash-frozen and stored at −80 °C until RNA isolation.

### RNA extraction, library preparation, and sequencing

Total nuclear RNA was extracted using the TRIzol LS protocol and resuspended in nuclease-free water. RNA concentration was determined using a Qubit RNA HS assay (Thermo Fisher, Q33221), and RNA integrity was assessed using a Fragment Analyzer (Agilent). PolyA^+^ RNA was isolated with the NEB-Next Poly(A) mRNA Magnetic Isolation Module (NEB, E7490L), and unique dual-indexed libraries were prepared with the NEB-Next Ultra II non-directional RNA Library Prep Kit (NEB, E7770L) with 17 cycles of PCR amplification. Because nuclear RNA is enriched for incompletely spliced pre-mRNA, polyA^+^ selection captures both mature and nascent polyadenylated transcripts. Libraries were quantified by Qubit (dsDNA HS) and sized on a Fragment Analyzer. Sequencing was performed on an Illumina NovaSeq X (2 × 150 nt paired-end) to a target depth of ≥20 M read pairs per library. A total of 13 libraries were sequenced (*Vglut2*^Cre/+^;*Sun1-sfGFP^1^*^x/+^: 4 fed, 3 fasted ; *Vgat*^Cre/+^; *Sun1-sfGFP*^1x/+^: 3 fed, 3 fasted).

### Quantification and statistical analysis

#### Library alignment, quality checks and expression analysis

Reads were aligned to GRCm38/mm10 with STAR v 2.7.0e ^27^. The following parameters were used for mapping: --outSAMstrandField intronMotif, --outFilterIntronMotifs RemoveNoncanonical, --outSAMtype BAM SortedByCoordinate, --quantMode GeneCounts. Two quantifications derive from this pipeline: the differential expression analyses underlying the volcano and response-magnitude panels (Figures 2G, 4B, 5B) used the original per-sample gene-body quantification, whereas all other analyses used the cleaned quantification. Both derive from the identical STAR/gene-body pipeline and differ only in sample inclusion and QC. Prior to normalization, per-gene counts were capped at 10,000 to limit the influence of highly abundant nuclear non-coding transcripts on size-factor estimation. Size factors were then estimated by the DESeq2 median-of-ratios method, and genes with a DESeq2 baseMean < 10 were excluded from differential testing. Counts per million (robust FPM) were used for baseline expression comparisons, and the variance-stabilizing transformation (VST) for principal-component analysis.

Differential expression was performed with DESeq2^28^ separately within each cell type, modeling counts with the design ∼ condition and testing the effect of fasting with Wald test statistics (24-hour fasted vs. ad libitum fed). Dispersions were estimated by DESeq2’s empirical-Bayes shrinkage across genes; this genome-wide dispersion estimation yields stable per-gene variance estimates at the replicate numbers used (Vglut2^Cre/+^; Sun1-sfGFP^1x/+^, n = 4 fed / 3 fasted; Vgat^Cre/+^; Sun1-sfGFP^1x/+^, n = 3 fed / 3 fasted after outlier exclusion). Log₂ fold changes are reported as maximum likelihood estimates with 95% confidence intervals (estimate ± 1.96 × SE). Significance was determined from adjusted p-values (FDR) following the Benjamini–Hochberg (BH) procedure. For the genome-wide differential-expression panels (volcano plots; Figures 4B, 5B), adjusted p-values are the standard DESeq2 transcriptome-wide BH values, and the same transcriptome-wide BH values were used for the heatmap and forest plots depicting the curated gene panels for GABA_A_ receptor subunits, GABAergic synthesis and release machinery, and glutamatergic signaling (Figures 4C–4F, 5C–5D). Significance was reported at a primary threshold of FDR < 0.10, with FDR < 0.25 noted as a secondary, exploratory tier.

The genome-wide distributions of fasting log₂ fold changes for the two cell populations (Figure 2G) were compared using a two-sample Kolmogorov–Smirnov test, which assesses whether two empirical cumulative distribution functions differ in shape without distributional assumptions. The statistic D is the maximum absolute difference between the two cumulative distributions (D = 0.351; p < 2.2 × 10⁻¹⁶). This is reported as a descriptive comparison of overall response distributions, not a per-gene inference. For each curated functional gene class (Figure 2H), per-gene baseline bias was computed as the log₂ ratio of mean ad libitum-fed CPM between populations (log₂[Vgat/Vglut2]) and summarized as the class mean ± SEM across genes, with individual gene values indicated. The directional bias of each gene class was assessed by counting the genes that were higher in each population and evaluating the split using a binomial (sign) test. Finally, individual transcripts and their baseline expression between VGLUT2^+^ and VGAT^+^ neuronal populations under ad libitum-fed conditions (Figure 3) were classified as expressed or not expressed, with genes classified as detected if the mean CPM for samples from the fed condition exceeded 10 CPM.

#### Gene annotations

Genes were tracked by NCBI Entrez ID; symbols were joined post hoc via org.Mm.eg.db. Analyses used R-studio with DESeq2^28^. Figures used ggplot2/patchwork (reversed magma heatmaps; viridisLite) and were assembled in Adobe Illustrator.

#### Quantification of Gabrd/Slc17a6 co-expression (Figure 5E)

DAPI^+^ nuclei within the P2rX2-defined rNTS were segmented, and a cell was scored as probe-positive if it contained at least 4 puncta within a 10-15 µm radius of the nucleus, a threshold set relative to the DAPI-negative control signal. The percentage of *Gabrd^+^* cells (as a fraction of DAPI^+^ cells) was compared between *Vglut2^+^* and *Vglut2⁻* populations (n = 3 mice). Quantification was performed by an experimenter blinded to genotype and condition.

#### Statistical analysis

Summary data are presented as mean ± SEM unless otherwise stated. The specific test, statistic, and n for each panel are given in the corresponding Figure legend. All analyses were performed in RStudio (v2025.05.1+513).

#### Software and figure preparation

Figures were generated in R using ggplot2 Version 2.0.0. Heatmaps use reversed magma from viridisLite and were assembled in Adobe Illustrator 2026.

## Notes

### Competing Interest Statement

The authors have declared no competing interest.

